# Kinship, acquired and inherited status, and population structure at the Early Bronze Age Mokrin necropolis in northern Serbia

**DOI:** 10.1101/2020.05.18.101337

**Authors:** Aleksandra Žegarac, Laura Winkelbach, Jens Blöcher, Yoan Diekmann, Marija Krečković Gavrilović, Marko Porčić, Biljana Stojković, Lidija Milašinović, Mona Schreiber, Daniel Wegmann, Krishna R. Veeramah, Sofija Stefanović, Joachim Burger

**Author notes:** corresponding author (A.Ž.), (J.Bu.).

## Abstract

Twenty-four ancient genomes with an average sequencing coverage of 0.85±0.25 X were produced from the Mokrin necropolis, an Early Bronze Age (2,100-1,800 BC) Maros culture site in Serbia, to provide unambiguous identification of biological sex, population structure, and genetic kinship between individuals. Of the 24 investigated individuals, 15 were involved in kinship relationships of varying degrees, including 3 parent-offspring relationships. All observed parent-offspring pairs were mother and son. In addition to the absence of biological daughters, we observed a number of young women and girls with no biological relatives in our sample. These observations, together with the high mitochondrial diversity in our sample, are consistent with the practice of female exogamy in the population served by Mokrin. However, moderate-to-high Y-chromosomal diversity suggests a degree of male mobility greater than that expected under strict patrilocality. Individual status differences at Mokrin, as indicated by grave goods, support the inference that females could inherit status, but could not transmit status to all their sons. The case of a son whose grave good richness outstrips that of his biological mother suggests that sons had the possibility to acquire status during their lifetimes. The Mokrin sample resembles a genetically unstructured population, suggesting that the community’s social hierarchies were not accompanied by strict marriage barriers.

## INTRODUCTION

### Kinship studies in the reconstruction of prehistoric social structure

An understanding of the social organization of past societies is crucial to understanding recent human evolution, and several generations of archaeologists and anthropologists have worked to develop a suite of methods, both scientific and conceptual, for detecting social conditions in the archaeological record (1–4). These methods have been used to investigate when social complexity, including social inequality, first appeared (5–8), the nature and function of early forms of social stratification, and how these emergent structures were perpetuated over time and space (9–11).

In the absence of written records, prehistoric social structure is reconstructed primarily via evidence from mortuary remains (12). Archaeological kinship studies use mortuary evidence to understand the specific role of family structure in shaping social organization (13, 14), and are critical for determining how familial relationships have influenced the emergence of social complexity and the evolution of persistent inequality (5, 10, 15, 16).

The anthropological understanding of kinship embraces not only biological relatedness, but also a broad range of non-biological social relationships (12). Recently, ancient DNA (aDNA) has been developed as a complementary line of evidence for reconstructing prehistoric kinship ties. Ancient DNA directly illuminates only one facet of past societies: biological relatedness. However, when combined with multiple lines of bioarchaeological and archaeological evidence, palaeogenetic data enable a comprehensive exploration of the family concept and social organization (17–21).

### The emergence of vertical differentiation in the Early Bronze Age

There is little evidence for significant social stratification before the end of the Pleistocene (7). It was the stable climatic conditions of the Holocene that enabled the adoption of sedentism and plant and animal domestication, stimulating massive increases in population growth (22). This growth, in turn, affected the economic structure of prehistoric societies, eventually leading to even greater population pressure and changes in social organization (7, 23). In the Early Bronze Age (EBA) we begin to detect pronounced levels of social inequality expressed through material culture and burial customs (24, 25). Economic innovations such as the long-distance exchange of ideas, knowledge, and “exotic” goods enabled significantly greater accumulation of material wealth, as well as the differential control over valuable resources. This increased social and economic complexity supported the emergence of elite individuals (25–27). EBA social complexity may have had biological relevance: in ranked societies, social status influences access to resources and positions of power, potentially of crucial importance for individuals’ health and fertility (5, 10).

With the establishment of unambiguous evidence for social inequality in the EBA, a debate has arisen surrounding the mechanisms perpetuating status and wealth. It is hypothesized that in complex societies such as Bronze Age chiefdoms, kinship networks and affinal ties served as conduits through which the leadership distributed wealth, controlled labor, and reinforced its own status (25, 27). Lineage-based intergenerational transmission of wealth is believed to have been the main mechanism for persistence of wealth inequality (7, 23).

### The role of Mokrin in understanding EBA social organization

Here we report the findings of a study combining palaeogenomic, bioarchaeological, and anthropological evidence to conduct a kinship analysis of the EBA necropolis of Mokrin, located near the town of Kikinda in the northern Banat, Serbia. The necropolis was used by a population belonging to the Maros culture (2700-1500 cal BC), which encompasses a set of communities extending through southeastern Hungary, western Romania, and northern Serbia ((28); Fig. 1; SI 1). Radiocarbon dating indicates that the Mokrin necropolis was used for 300 years, from around 2100-1800 cal BC (29), Table S1). Large and well-preserved cemeteries are typical of Maros settlements, and Mokrin, with a total of more than 300 graves, is one of the largest. The siting of Maros villages along rivers and the presence of distinctively non-local grave goods indicate that Maros communities engaged in frequent interactions with non-local groups, including the exchange of manufactured goods and resources (30). The bioarchaeological and material-culture correlates of social organization at Mokrin have been well-characterized through detailed analyses of the variability and spatial distribution of grave goods (30), (SI 2); and the ways in which physical activity patterns, discerned from skeletons, reflect wealth, status, and socio-political factors (31). These lines of research have identified grave goods that functioned as markers of higher social status (Fig. S1) (30), and detected a positive correlation between these markers and increased male physical activity (31, 32).

**Fig. 1.**
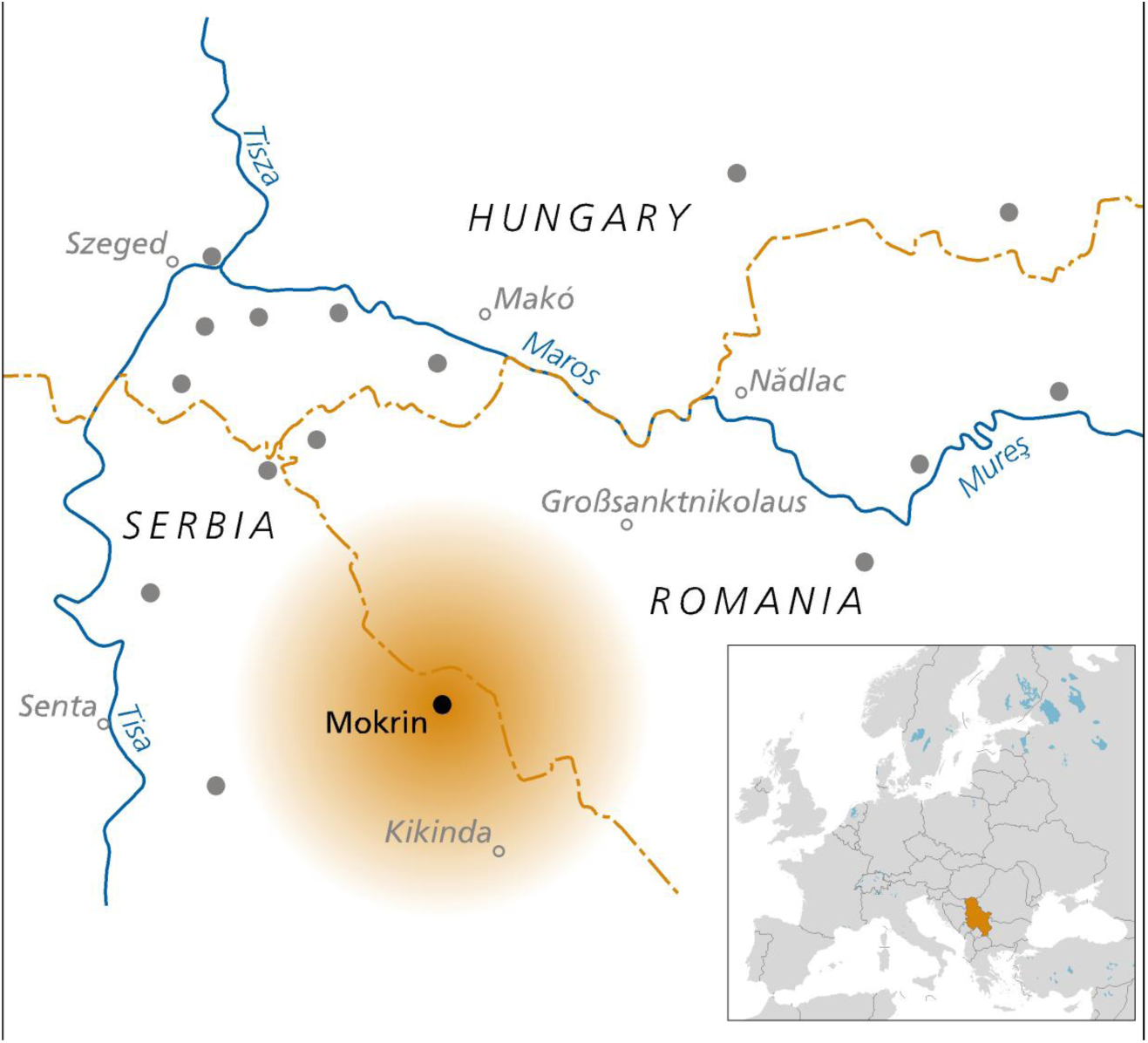
Southeast Europe - the Mokrin necropolis Dots represent other known Maros settlements and cemeteries.

In this study, we conducted palaeogenomic analyses on 24 skeletal samples from the Mokrin necropolis to provide unambiguous identification of biological relatedness between individuals. Identified kinship ties are used together with archaeological markers of social status to infer features of developing vertical differentiation within EBA Maros society, and— more broadly—to trace aspects of the evolution of Bronze Age societies.

We address four main questions:

1. What was the kinship system at the community served by the Mokrin burial site: were families organized in clans, lineages, or larger kindreds, and can we reconstruct residence and marital patterns?
2. Were wealth and status hereditary in the society represented by the Mokrin assemblage?
3. Does the genetic variability in the Mokrin sample correspond to that of a single population? Is there evidence of inbreeding?
4. How is the Mokrin assemblage representative of the genetic diversity during the Copper Age–Bronze Age transition?

## RESULTS

### Sampling and anthropological analysis

We selected 24 individuals (14 adults and 10 children) buried in 22 graves for analysis according to a suite of criteria including petrous bone preservation, the presence of neighbouring graves, the presence of younger individuals buried in close proximity to adults (as potential family groupings to track the inheritance of status), and variety in material culture markers (Table 1, Fig. 2, SI 2). We primarily sampled single graves except for the double-burial 257 and triple-burial 122 (Fig. S2). Sixteen out of 24 skeletal remains were taphonomically well preserved (SI 1, Table S2). Palaeopathological and physiological stress markers were typical for a prehistoric population of this area and period (See SI 1 for more details).

**Table 1.** Early Bronze Age Mokrin samples analyzed: burial information, genetic sex determination, age, grave goods, genomic sequencing coverage and haplotype information.

| Burial | Genetic Sex | Age | Grave goods | X-fold genomic depth | mt haplogr. | Y haplogr. |
| --- | --- | --- | --- | --- | --- | --- |
| 122E | XY | 6-9 | bone needle, kaolin and bone bead necklace, bronze earring | 1.09 | U5a2b1a | I2a1b |
| 122S | XX | 35-50 | *Columbella* shells, *Dentalium*, animal teeth and kaolin necklace, bronze bracelet, bronze head ornament | 0.78 | H32 | * |
| 161 | XX | 9-11 | necklace consisting of kaolin and *Dentalium* beads, and animal bones and teeth, bronze head ornament, bone needle, bronze ring | 1.20 | H80 | * |
| 163 | XY | 45-55 | biconical vessel, stone hammer-ax, biconical beaker | 1.21 | U4a2 | J2b |
| 181 | XX | >18 | biconical beaker, necklace made of *Dentalium* snail, deer tooth, kaolin and an oblong plaque of copper sheet | 0.62 | U4a2 | * |
| 186 | XX | 8-11 | biconical beaker, biconical vessel | 0.33 | H1aj | * |
| 211 | XY | 50-55 | bigger bowl, bronze dagger | 0.79 | U5a2b1a | I2a1b |
| 220 | XY | 15-25 | - | 0.64 | T2b11 | R1b1a2a2c1 |
| 223 | XX | 7-10 | beaker<br>biconical vessel | 0.39 | U3a1 | * |
| 224 | XX | 25-40 | - | 0.77 | T2b | * |
| 225 | XY | 25-35 | - | 0.82 | J1b1a1 | R1b1a2a2c1 |
| 228 | XX | 35-50 | ball-like beaker, beaded sash (bone pearls, a bead made of *Dentalium* shell, a kaolin clay spindle whorl) | 0.95 | J1c | * |
| 237 | XX | 15-20 | bronze head ornament, kaolin and *Dentalium* necklace, bronze bracelet, biconical bowl, biconical amphora | 0.89 | T2b | * |
| 243 | XY | 20-35 | biconical beaker, stone ax | 1.12 | H | BT |
| 246 | XX | 45-50 | Amphora, lid, head ornament consisted of bronze plaques, *Columbella* shell and pendants, necklace made of bronze bead, kaolin and *Dentalium* snail, Unionidae freshwater mussel, two bone needles | 0.98 | H80 | * |
| 247 | XX | 10-12 | kaolin, *Dentalium* and animal teeth necklace, bone needle, sheep jaw | 0.90 | H1 | * |
| 257 A | XX | 40-60 | - | 0.60 | H | * |
| 257 B | XY | inf. I | biconical beaker | 0.61 | K1a4 | R1b1a2a2c1a1 |
| 260 | XY | 15-18 | - | 0.92 | J1c | I2a2a1a2a2 |
| 282 | XY | 15-20 | biconical beaker, bellied beaker with specific pattern, biconical bowl | 1.41 | H2b | BT |
| 287 | XX | 20-35 | bronze head ornament, copper chisel (possibly used for trepanation), two bone needles, copper bracelet, necklace made of *Columbella* shell and *Dentalium*, copper and kaolin beads, gold pendant, two biconical vessels | 0.81 | U5b2a2c | * |
| 288 | XX | 60+ | two copper bracelets, necklace made of *Dentalium* and kaolin beads, two bone needles, biconical bowl, biconical beaker | 0.81 | HV0e | * |
| 295 | XY | 15-20 | cylindrical cup, biconical bowl | 0.82 | H80 | I2a1a |
| 302 | XX | 20-35 | Beaker, conical bowl, copper head, ornament with kaolin, animal teeth and bones, *Dentalium* beads, *Columbella* shell and *Cardium* sp. | 0.89 | J1c | * |

**Fig. 2.**
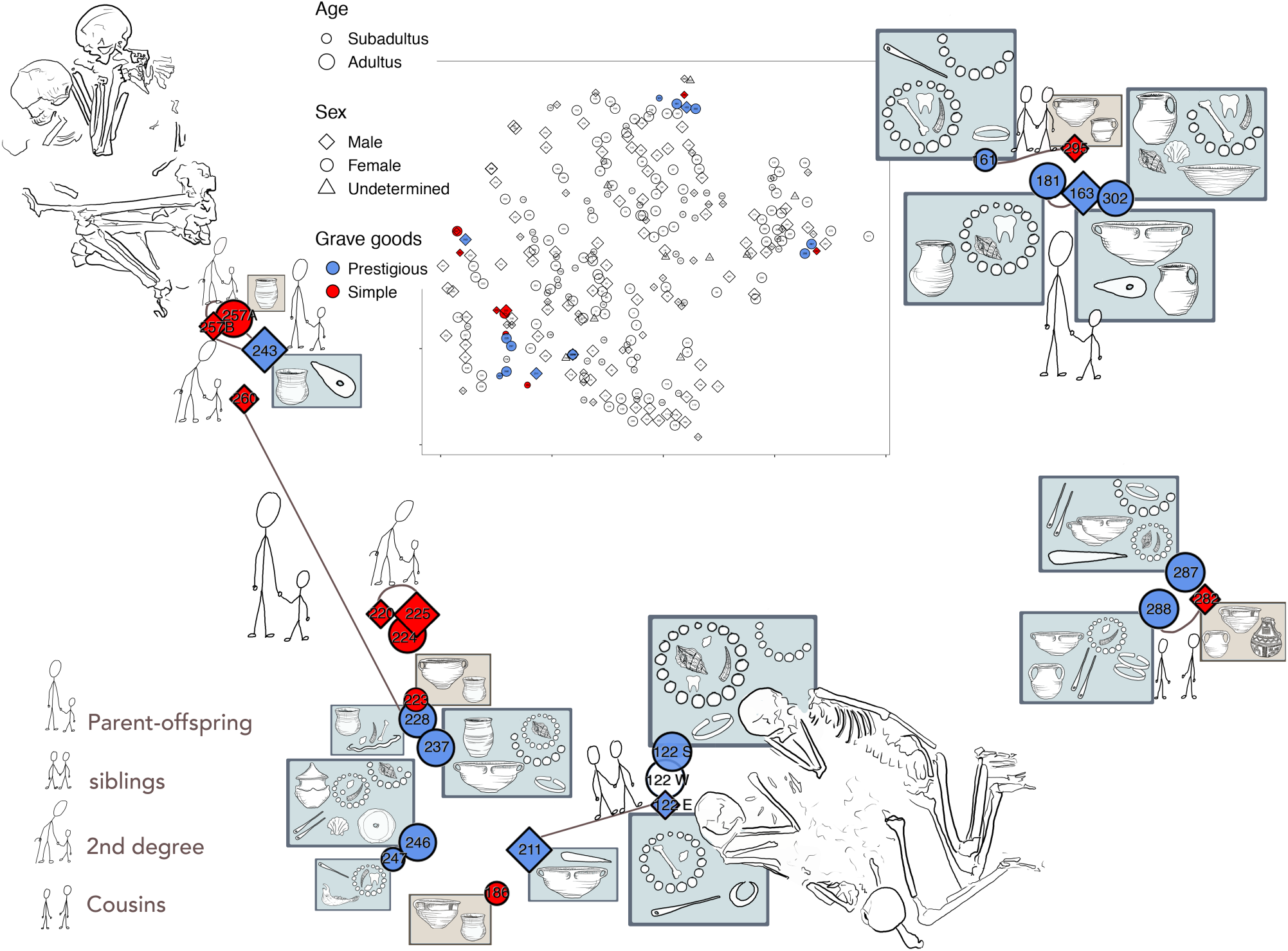
Visual representation of the 24 analyzed individuals and their relationships of the Mokrin necropolis. Head ornaments are represented by semi-circles, necklaces by closed circles, and beaded sashes by wavy lines. The material used for making the jewelry is indicated by the symbol inside the jewelry.

We divided our sample into two categories according to the richness of their grave goods: ‘prestigious’ (characterized by numerous and status-indicating goods in their burials) and ‘poor’ burials (those having few or simple grave goods or none at all) (30).

### Assessment of DNA-preservation, contamination, post-mortem damage, and sex

We extracted DNA from petrous bones and performed whole genome shotgun sequencing, reaching an average depth of 0.85X±0.25 (0.33X - 1.41X) on the autosomes (Table 1). The proportion of endogenous human DNA in the 24 investigated skeletons ranged from 8% – 70% (with only two samples below 20%), reflecting the very good molecular preservation of these samples (SI 3, Table S5). Excellent preservation was also indicated by the low levels of contamination detected in the samples: the average mitochondrial contamination level was < 1% as estimated by contamMix (Table S5) (33). For non-USER treated libraries, deamination rates ranged from 0.13 to 0.26 at the first base of the 5’ end of the reads (Table S5, SI 3), supporting the authenticity of the aDNA data.

Fourteen females and ten males were identified via molecular sexing (34); (Table 1), confirming the anthropological sexing of the remains. We observed only three discrepancies in molecular and anthropological sex assignment, all in individuals whose sex was initially determined according to the funerary ritual alone (122S, 220 and 257B) (28).

### Uniparental markers and genomic diversity estimates

The Mokrin sample displays relatively high haplogroup diversity for both the non-recombining portion of the Y-chromosome (NRY, h=0.81) and mitochondrial DNA (mtDNA, h=0.95) (Table 1). We discerned at least 14 distinct mtDNA haplotypes, including several belonging to haplogroup U, often found in prehistoric Central European foragers (35), as well as to H, T2, K1, and J1. The ten Y chromosomes could be assigned to at least five different haplotypes, of which three belonged to the R1b family common among modern European populations (36). No evidence of significant population genetic structure was found. We estimated the inbreeding coefficient *F* (defined as a deficit of heterozygote genotypes given population allele frequencies) to be zero, following the method described in (37). We additionally tested several hypothesized partitions of our sample (SI 7), but did not observe significant population differentiation in any configuration (F_ST_ ≈ 0).

### Ancestry analyses

When projected onto a PCA of European populations, all Mokrin samples fall within modern European genetic variation, clustering in the midst of modern northern, eastern, and southern Europeans (Fig. S5).

We estimated individual admixture proportions under the assumption that the composition of a European Bronze Age population can be sufficiently modeled with three components: western hunter gatherers, Aegean Neolithic farmers, and eastern European steppe-like populations. We observed no significant variation in the eastern European steppe-like component between individuals (Fig. S6, Table S6). Pooling individuals, admixture proportions are estimated to be around 8% (± 1.2% standard error (SE)) western hunter gatherers, 55% (± 2.5% SE) Aegean Neolithic farmers, and 37% (± 2.3% SE) Eastern European steppe-like population (Fig. S7). Quantification of shared drift to other temporally and geographically close ancient individuals via outgroup f3 statistics did not reveal any particularly close affinities (Fig. S8), reflecting the genetic homogenization of Europe during the Bronze Age.

### Biological relatedness analysis

Genome-wide patterns of identity-by-descent among the 24 analyzed individuals from the Mokrin necropolis revealed nine family relationships involving 15 Mokrin individuals (Table 2, Fig. 2). In addition to three parent-offspring and two sibling relationships, we reconstructed three second-degree (half-siblings, avuncular, grandparent-grandchild) and one third-degree (cousin) relationship. Inferred kinship relations were corroborated by an outgroup f_3_ analysis in which related individuals clustered together tightly (Fig. S8). Related individuals tended to be buried in close proximity (permutation test, p < 8.5 * 10^−4^), with two exceptions (Fig. 2). Nine out of 24 individuals did not have a close genetic relationship to any other individual (186, 122S, 223, 224, 237, 246, 247, 287, 302); they were all female (3 young girls and 6 adult women, permutation test p < 1.6 * 10^−3^). These women are evenly distributed over the entire sampling area.

**Table 2:** Results of the kinship analysis obtained by lcMLkin software.

| Biological relationship | Individual 1 |  | Individual 2 |  |
| --- | --- | --- | --- | --- |
|  | Burial | Sex and age | Burial | Sex and age |
| Parent-offspring relationship |  |  |  |  |
|  | 257A | ♀, adultus | 243 | ♂, adultus |
|  | 228 | ♀, maturus | 260 | ♂, juvenis |
|  | 163 | ♂, adultus | 181 | ♀, adultus |
| Sibling relationship |  |  |  |  |
|  | 122E | ♂, infans II | 211 | ♂, maturus |
|  | 161 | ♀, infans II | 295 | ♂, juvenis |
| Second degree relationship (half-siblings, avuncular, grandparent-grandchild) |  |  |  |  |
|  | 257B | ♂, infans I | 257A | ♀, adultus |
|  | 257B | ♂, infans I | 243 | ♂, adultus |
|  | 220 | ♂, juvenis | 225 | ♂, adultus |
| Cousin relationship |  |  |  |  |
|  | 282 | ♂, juvenis | 288 | ♀, senior |

### Phenotypic markers

We estimated frequencies of a set of markers related to pigmentation phenotype in the Mokrin sample by calculating individual genotype likelihoods using a Bayesian approach implemented in ATLAS (38). The frequency of the derived allele at rs16891982*G (*SLC45A2*) was 0.7098 (CI 0.5365 - 0.8476; N=15) and at rs1426654*A (*SLC24A5*) 1 (CI 0.8899 - 1; N=15); both are associated with skin depigmentation in Europeans (39). Comparable frequencies can be found in modern day populations in Spain (*SLC45A2*: 0.8178, *SLC24A5*: 1). The frequency of the derived G allele at rs12913832 in the *HERC2* gene, which is strongly associated with iris depigmentation, was estimated to be 0.4498 (CI 0.2946 - 0.6127; N=20), similar to modern day populations in Tuscany (0.4206).

## DISCUSSION

### The Mokrin population: ancestry, structure, and genetic diversity

The individual Mokrin genomes are best modelled as a mixture of Central European hunter-gatherers, Aegean Neolithic farmers and influences from the Eastern European steppes (mean *qpAdm* tail probability individually 0.46, pooled 0.08). The Aegean/Mediterranean ancestry component dominates in our sample (pooled 55% ± 2.5%), while the hunter-gatherer component is relatively low (pooled 8% ± 1.2%), and indeed it is statistically undetectable in four individuals.

The estimated inbreeding coefficient (*F*) is very low, suggesting that the necropolis represents a randomly-mating population. We found a rather high number of mitochondrial lineages (14 haplotypes in 24 individuals). High mtDNA diversity in combination with archaeological and isotope evidence can indicate female exogamy (21, 40, 41). Given the absence of genetic substructure and the fact that both Y-chromosomal and mitochondrial diversity is moderate-to-high at Mokrin, the most parsimonious explanation is that this cemetery served a single, large, and contiguous population.

### Biological relatedness and implications for the Mokrin kinship system

Genetic kinship analysis, which revealed close genetic relationships involving 15 of the 24 burials analyzed, provided insight into the mortuary practices at the Mokrin site.

Our kinship analysis did not identify any fathers or daughters, and consequently, there were no mother-father-child trios. We did not find genetic evidence for larger kindred, extended families, clans or lineages in our sample: this could be due to the limited number of samples analyzed (24 out of 312 excavated burials), or insufficient power to reliably detect genetic relationships beyond the 3^rd^ degree. We therefore cannot exclude the possibility that larger family units were buried scattered at the cemetery.

Interestingly, all nine of the women without genetic relatives in our sample were found in close proximity to other burials. The adult woman 122S, for example, was buried in the only triple burial of the entire necropolis. Double and triple burials are highly unusual in Maros cemeteries, and they imply special burial circumstances such as a traumatic event resulting in the death of several people in quick succession and their joint interment (SI 2). Two of the “unrelated” females could be assumed, based on archaeological evidence, to be the wives of two men buried nearby, but the family relationships of the other seven women are unclear. The locations of their burials, however, hint that they might not have been entirely socially isolated in the local community. The absence of detected biological daughters and the presence of women of different status with no kin in the cemetery, considered together with the high mtDNA diversity observed in our sample, suggests that female exogamy may have been practiced in Mokrin’s source population.

While the above observations do not support an inference of strict matrilocality, we cannot definitively conclude that the Mokrin society was patrilineal and/or patrilocal. The observation that the necropolis was used by a single unstructured population in combination with a moderate to high Y-chromosomal haplotype diversity (h=0.81) militates against strict patrilocality, as we then would expect to see a comparatively lower Y-chromosomal diversity.

### Patterns of status transmission at Mokrin

Our sampling strategy targeted burials located close to each other within the Mokrin complex on the assumption that these would be more likely to contain genetically related individuals. Unraveling these biological ties through kinship analysis enabled us to infer whether wealth and social status (as indicated by grave goods) were inherited or earned in the Maros society represented by the Mokrin assemblage. If social prestige was transmitted intergenerationally, we would expect to see this reflected in status markers. Based on previous palaeogenomic and isotopic research from a similar period of prehistory (21, 41), we hypothesized that social status would be transmitted along patrilines, with women acquiring their status through marriage.

Although the differences in status within the Mokrin necropolis were not extreme, there was great variability in grave goods displayed by biological relatives.

There were only two cases in which relatives expressed similar social status through their grave goods: the two men in burials 220 (15-25 years old) and 225 (25-35 years old) were second degree relatives and were both buried without grave goods. At the opposite end of the status spectrum, a woman from the burial 181 was very likely the mother of the male in burial 163 and both were buried with grave goods indicative of higher social status (30). Since the male individual was an adult at the time of death, it remains unclear whether he inherited or acquired his status.

Other observations at Mokrin do not support the inference that social status was transmitted intergenerationally to males. For example, the woman in burial 228 was the mother of the subadult male in burial 260 (15-18 years old). While the mother had comparatively rich grave goods, her son was buried without any grave goods at all. We infer that this subadult neither inherited his mother’s social status nor acquired it in the course of his young life. Similarly, the woman 257A was identified as the mother of the 20-35 years old male buried in grave 243. Her grave goods suggest that she was of lower status, while her son’s grave contained an ax, an indicator of higher status. This status discrepancy suggests that the son acquired the status to command a richer burial during his life. We additionally note that the contrast in grave good richness observed in this quartet of burials is not consistent with the inheritance of status via the maternal line.

Evidence from another pair of burials demonstrates that sub-adults could have rich grave goods—if they were girls. A burial of a 9-11 years old girl (burial 161) contained various markers of higher status (a necklace, bronze head ornament, a bone needle, and a bronze ring). Her brother (burial 295), who was 15-20 years old when he died, had only small and simple ceramics beside him. The fact that these siblings display different status in the grave is more consistent with a system wherein either females—but not males—could inherit social status. This case is particularly convincing evidence that only women could inherit status, as the girl was almost certainly too young to have had the opportunity to acquire status, and must have inherited her rich adornments. However, an alternative explanation is that only very young children—but not teenagers/adolescents—inherited status in the grave. We further observed a cousin relationship between a higher-status adult female (288) and a lower-status juvenile male (282). The dissimilarity in status between these cousins is consistent with the idea that males did not inherit status, but does not rule out a system in which male children were assigned the status of their fathers until they reached a certain age or level of merit.

In total, we inferred biological kinship relationships for ten males in our sample. Given the observed status and age distribution, we can rule out that the higher–lower social status dichotomy in males is exclusively due to age differences in our sample. When we consider only first-degree relationships, for three out of six males it seems unlikely that status was inherited (burials 243, containing a 20-25 years old; 260, 15-18 years old; and 295, 15-20 years old), as their immediate female relatives (mother or sister) differed from them in their grave good status. For two other males (163, age 45-55; and 211, age 50-55), it is unclear whether status was inherited or acquired. The most problematic case for our interpretation is burial 122E, a 6-9 year-old boy buried with markers of higher social status—seemingly clear evidence of inherited status. However, 122E was part of a triple burial and the assignment of the grave goods to the boy is not completely secure.

Taken together, these observations do not support the inference of inheritance of social status in men; male status appears to be acquired, except for the inconclusive case of the boy in grave 122. However, our sample does not include any of the fathers of the buried men and boys. We must therefore limit the scope of our claim to posit that sons in our sample have not inherited social status from their mothers.

### Reconstructing social organization at Mokrin/Mokrin in its temporal context

This analysis of 24 ancient genomes has illuminated important features of the social organization of the Early Bronze Age society served by the Mokrin necropolis, particularly concerning the inheritance of status.

As already mentioned, the Mokrin skeletal sample appears to represent a genetically unstructured population. While this does not exclude the existence of social hierarchies, it does indicate that there were no strict barriers to marriage between social groups.

Multiple lines of evidence in our sample—high mtDNA variability, the presence of a certain number of unrelated women, and the absence of daughters—indicate that female exogamy was practiced between Mokrin and other settlements. Interestingly, the absence of adult daughters and presence of unrelated females has also been reported at a Bronze Age settlement in Bavaria (21). However, unlike the unrelated Bavarian females, who were mostly of higher status, the unrelated Mokrin females display a wide range of grave good richness, from poor to prestigious.

At the Mokrin necropolis, relatives were buried close together in small kinship groups; interestingly, in our sample these small groups did not include biological fathers. The absence of larger kindreds and the relatively high NYR diversity in our sample are evidence against strict patrilocality in this population. These observations suggest a different form of social organization from that of a Bell Beaker group in southern Germany, an assemblage containing high mtDNA variability but only a single Y-chromosomal lineage. However, the low Y-variability here could be typical for the entire region at this time and have no social implications at all (41).

Status inheritance at Mokrin also appears to have differed from other EBA cultures. It seems that sons did not inherit social status from their biological mothers, but had the opportunity to acquire status throughout their lives. It is possible that sons may have inherited their status from their fathers, but this would require that spouses display different status in the grave. An alternative explanation is that in Mokrin the law of the first-born was valid and in our sample only the post-born sons are present. In any case, the situation in Mokrin is different from the EBA Lech valley and the Bell Beaker population in Bavaria where clear signals of (male) status inheritance are observed.

Our kinship analysis identified three mothers, one sister, and nine unrelated females, but no daughters, complicating inference about inherited or acquired status in females. Here Mokrin once again differs from the Bell Beaker sites in southern Germany, where girls are under-represented in burial assemblages. In Mokrin, the sex-dependent body orientation is almost universally applied to adults and children of both sexes, an inhumation pattern which is in accordance with the highly normative funerary ritual observed in all Maros cemeteries.

It is evident from the few existing palaeogenetic studies on this topic that there is significant regional and temporal variability in the social structures and heredity patterns of Late Neolithic and EBA societies, although one common thread among the different societies investigated appears to be the practice of female exogamy. By illuminating the development of vertical differentiation and heredity systems, complete analyses of large cemeteries like Mokrin can help to trace the evolution of Early Bronze Age societies.

## METHODS

### Production of palaeogenomes

Sample preparation of petrous bones from 24 individuals was carried out in the aDNA laboratories of the Palaeogenetics group at the Johannes Gutenberg University Mainz following the established protocol described in SI 3. Double-indexed libraries were prepared according to (42) with modifications (see SI 3) and screened on an Illumina MiSeq™ platform at StarSEQ GmbH (Mainz, Germany). For deeper shotgun sequencing, aDNA extracts were treated with USER™ enzyme (43) prior to library preparation. Whole-genome sequencing was performed on Illumina’s NovaSeq™ 6000 platform at the Next Generation Sequencing Platform (Institute of Genetics) at the University of Bern (Switzerland). For details see SI 3.

### Bioinformatic analyses

Raw data was analyzed with bioinformatics methods adjusted to account for the unique properties of aDNA, as described in detail in SI 4. Reads were aligned against the reference genome (GRCh37/hg19) using bwa aln (44). During the conversion to the BAM format, reads were filtered for a minimal mapping quality of 30. PCR duplicates were marked using sambamba (45) prior to realignment with GATK (46) around known SNPs and InDels. Authenticity of DNA was assessed based on the mitochondrial chromosome using ContaMix (33) and postmortem damage patterns (deamination at 5’ and 3’ ends) were quantified with MapDamage2 in aligned sequence reads for non-USER treated libraries (47).

SNP calling was carried out following the approach described in (48) using the ATLAS package (38). Genotype calls were obtained with the maximum likelihood approach described in (48). Majority-allele calls were produced for the SNPs overlapping the 124k capture array described in (49), the Y-chromosome and the mitochondrial chromosome. In each case, sequencing errors and post-mortem damage were considered during variant detection. In addition, genotype likelihoods were calculated per individual and used for allele frequency estimates.

The molecular sex of each sample was determined following the approach described in (34). Y-chromosomal haplotypes were predicted with the yHaplo tool (50), while mt-haplotypes were determined using Haplogrep 2.0 (51).

### Affinities and Ancestry

For inferences about genetic affinities and ancestry, we computed a PCA with LASER (52) against a reference space of modern European individuals (53) and f-statistics with *qp3Pop, qpDstat* in *f*_*4*_ mode and *qpAdm* from the ADMIXTOOLS package (54) (SI 5). *qpAdm* standard errors are computed by jackknifing excluding successive 5cM blocks.

### Biological relatedness

Biological relatedness within the cemetery was estimated with lcMLkin (55). The underlying assumption of these analyses is that close genetic relatives are more similar due to sharing alleles that are identical by descent because they were inherited from a recent common ancestor. Given the above genetic background, we performed genotype calling on the Mokrin samples at 6,191,202 SNPs that have a frequency of >=5% in the 1000 Genomes Eurasian samples (56). We then performed biological relatedness estimation using three approaches on this set of SNPs (SI 6). All results were confirmed using pairwise distances and READ software using the original set of 6,000,000 SNPs, though READ was only able to identify relationships up to the second degree (SI 6). We assessed whether relatives tend to be buried close to each other using a permutation test. For this, we grouped the individuals into the four geographic groups north-east (287, 288, 282), north-west (257A, 257B, 243, 260), south-east (161, 295, 181, 163, 302) and south-west (220, 225, 224, 223, 228, 237, 246, 247, 186, 211, 122S, 122W, 122E) and quantified the number of pairwise relationships in which both individuals were buried in the same group. To assess significance, the observed number (eight) was compared against those obtained in 10^7^ permutations of the individuals among groups.

### Population genetic diversity estimates

The full sets of individuals we hypothesised may constitute clusters in a structured population are northern (161, 163, 181, 302, 287, 288, 243, 257A, 257B, 295, 260, 282) and southern (122E, 112S, 246, 247, 186, 211, 237, 220, 223, 224, 225, 228), individuals with (122S, 186, 223, 224, 237, 246, 247, 287, 302) and without (257A, 122E, 161, 163, 181, 211, 220, 225, 228, 243, 257B, 260, 282, 288, 295) family members buried in the necropolis, and an upper (181, 220, 287, 224, 225, 260, 228) and lower (all remaining) cluster in the PCA.

Pairwise distances were computed with PLINK (58) (1-IBS distance) based on autosomal sites with one or two alleles called with ATLAS maximum likelihood caller (38) as described above. The sets of individuals were filtered for relatedness, keeping only the genome with highest coverage in a family cluster of related individuals as given in Table 2, yielding the reduced set 122E, 122S, 163, 186, 220, 223, 224, 237, 243, 246, 247, 260, 282, 287, 295, 302. Comparisons between distributions of distances between groups were performed with a Kolmogorov-Smirnov Test as implemented in R.

We used the approach described in (37) to determine the inbreeding coefficient *F*. Using ATLAS, a multi-sample vcf file was generated containing the genotype likelihoods of each individual. F was estimated based on genome-wide SNPs with a minimum quality of 40 that were covered at least twice in a minimum of ten samples. MCMC was run for 10^6^ iterations with ten burn-ins of 500 iterations each. In addition we re-ran the analysis while restricting F>0.

Mitochondrial and Y-chromosomal haplotype diversity H was estimated as:

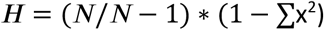

where x is the haplotype frequency, estimated among N samples.

### Allele frequencies of functional markers

Allele frequencies were estimated based on individual genotype likelihoods with ATLAS (38) using a Bayesian approach. Frequencies were compared to European population samples from the 1000 Genomes project (CEU: Utah residents with Northern and Western European ancestry, GBR: British from England and Scotland, IBS: Iberian populations in Spain, TSI: Toscani in Italia) by F_st_, using Hudson’s estimator (59).

## Acknowledgments

We would like to thank all of the colleagues for assistance and provided help in article preparation, especially to Mihailo Radinović, Dragoslav Stojanović, Ivana Živaljević and Tamara Blagojević. Parts of this research were conducted using the supercomputer Mogon and/or advisory services offered by Johannes Gutenberg University Mainz (hpc.uni-mainz.de).

## References

1. L. R. Binford, Mortuary Practices: Their Study and Their Potential. Memoirs of the Society for American Archaeology 25, 6–29 (1971).

2. C. Carr, Mortuary practices: Their social, philosophical-religious, circumstantial, and physical determinants. Journal of Archaeological Method and Theory 2, 105–200 (1995).

3. J. M. O’Shea, Mortuary variability: an archaeological investigation (Academic Press, London, UK, 1984).

4. C. S. Peebles, S. M. Kus, Some Archaeological Correlates of Ranked Societies. American Antiquity 42, 421–448 (1977).

5. K. M. Ames, “The Archaeology of Rank” in Handbook of Archaeological Theories., R. A. Bently, D. G. H. Maschner, C. Chippenale, Eds. (AltaMira Press, 2007), pp. 487–513.

6. G. M. Feinman, “The emergence of social complexity” in Cooperation and Collective Action: Archaeological Perspectives, D. M. Carballo, Ed. (University Press of Colorado, 2013), pp. 35–56.

7. S. M. Mattison, E. A. Smith, M. K. Shenk, E. E. Cochrane, The evolution of inequality. Evolutionary Anthropology: Issues, News, and Reviews 25, 184–199 (2016).

8. T. D. Price, G. M. Feinman, “Social Inequality and the Evolution of Human Social Organization” in Pathways to Power. Fundamental Issues in Archaeology, T. D. Price, G. M. Feinman, Eds. (Springer, New York, NY, 2010), pp. 1–14.

9. R. D. Drennan, C. E. Peterson, J. R. Fox, “Degrees and Kinds of Inequality” in Pathways to Power. Fundamental Issues in Archaeology, T. D. Price, G. M. Feinman, Eds. (Springer, New York, NY, 2010), pp. 45–76.

10. M. H. Fried, The evolution of political society: an essay in political anthropology (McGraw-Hill, New York: Random House, New York, 1967).

11. M. D. Sahlins, Stone age economics (Aldine - Atherton, Inc, Chicago, 1972).

12. K. M. Johnson, K. S. Paul, Bioarchaeology and Kinship: Integrating Theory, Social Relatedness, and Biology in Ancient Family Research. Journal of Archaeological Research 24, 75–123 (2016).

13. K. W. Alt, W. Vach, Odontologic kinship analysis in skeletal remains: concepts, methods, and results. Forensic Sci. Int. 74, 99–113 (1995).

14. C. M. Stojanowski, M. A. Schillaci, Phenotypic approaches for understanding patterns of intracemetery biological variation. Am. J. Phys. Anthropol. 131, 49–88 (2006).

15. K. M. Johnson, “Opening Up the Family Tree: Promoting More Diverse and Inclusive Studies of Family, Kinship, and Relatedness in Bioarchaeology” in Bioarchaeologists Speak Out, J. Buikstra, Ed. (Springer, New York, NY, 2019), pp. 201–230.

16. J. Müller, “Inheritance, population development and social identities: Southeast Europe 5200–4300 BCE” in Balkan Dialogues. Negotiating Identity between Prehistory and the Present, M. Gori, M. Ivanova, Eds. (Routledge, 2017), pp. 156–168.

17. W. Haak, et al., Ancient DNA, Strontium isotopes, and osteological analyses shed light on social and kinship organization of the Later Stone Age. Proc. Natl. Acad. Sci. U. S. A. 105, 18226–18231 (2008).

18. C. E. G. Amorim, et al., Understanding 6th-century barbarian social organization and migration through paleogenomics. Nat. Commun. 9, 3547 (2018).

19. C. Keyser-Tracqui, E. Crubézy, B. Ludes, Nuclear and mitochondrial DNA analysis of a 2,000-year-old necropolis in the Egyin Gol Valley of Mongolia. Am. J. Hum. Genet. 73, 247–260 (2003).

20. N. O’Sullivan, et al., Ancient genome-wide analyses infer kinship structure in an Early Medieval Alemannic graveyard. Science Advances 4, eaao1262 (2018).

21. A. Mittnik, et al., Kinship-based social inequality in Bronze Age Europe. Science 366, 731–734 (2019).

22. S. Shennan, The First Farmers of Europe: An Evolutionary Perspective (Cambridge University Press, 2018).

23. E. A. Smith, et al., Production Systems, Inheritance, and Inequality in Premodern Societies. Current Anthropology 51, 85–94 (2010).

24. T. K. Earle, Chiefdoms in Archaeological and Ethnohistorical Perspective. Annual Review of Anthropology 16, 279–308 (1987).

25. A. F. Harding, European Societies in the Bronze Age (Cambridge University Press, Cambridge, UK, 2000).

26. B. Hayden, S. Villeneuve, “Who Benefits from Complexity? A View from Futuna” in Pathways to Power. Fundamental Issues in Archaeology, T. D. Price, G. M. Feinman, Eds. (Springer, New York, NY, 2010), pp. 95–145.

27. K. Kristiansen, T. B. Larsson, The Rise of Bronze Age Society: Travels, Transmissions and Transformations (Cambridge University Press, Cambridge, UK, 2005).

28. M. Girić, Mokrin. Nekropola ranog bronzanog doba: Mokrin. The early bronze age necropolis (Dissertationes et monographie XI. Washington, Kikinda i Beograd: Smithsonian Institution, Narodni muzej, Arheološko društvo Jugoslavije, Jugoslavija, 1971).

29. J. M. O’Shea, A radiocarbon-based chronology for the Maros Group of southeast Hungary. Antiquity 66, 97–102 (1992).

30. J. M. O’Shea, Villagers of the Maros: A Portrait of an Early Bronze Age Society (Plenum Press, New York, NY, 1996).

31. M. Porčić, S. Stefanović, Physical activity and social status in Early Bronze Age society: The Mokrin necropolis. Journal of Anthropological Archaeology 28, 259–273 (2009).

32. S. Stefanović, Skeletal markers of occupational stress in later prehistory: Mokrin necropolis (2000–1800 B.C.) (University of Belgrade, Belgrade, Serbia, 2008).

33. Q. Fu, et al., The genetic history of Ice Age Europe. Nature 534, 200–205 (2016).

34. P. Skoglund, J. Storå, A. Götherström, M. Jakobsson, Accurate sex identification of ancient human remains using DNA shotgun sequencing. J. Archaeol. Sci. 40, 4477–4482 (2013).

35. B. Bramanti, et al., Genetic discontinuity between local hunter-gatherers and central Europe’s first farmers. Science 326, 137–140 (2009).

36. W. Haak, et al., Massive migration from the steppe was a source for Indo-European languages in Europe. Nature 522, 207–211 (2015).

37. J. Burger, et al., Genomic Data from an Ancient European Battlefield Indicates On-Going Strong Selection on a Genomic Region Associated with Lactase Persistence Over the Last 3,000 Years. CURRENT-BIOLOGY-D-20-00414, Available at SSRN: https://ssrn.com/abstract=3565013 or http://dx.doi.org/10.2139/ssrn.3565013 (submitted).

38. V. Link, et al., ATLAS: Analysis Tools for Low-depth and Ancient Samples. bioRxiv, 105346 (2017).

39. M. Soejima, Y. Koda, Population differences of two coding SNPs in pigmentation-related genes SLC24A5 and SLC45A2. Int. J. Legal Med. 121, 36–39 (2007).

40. H. Schroeder, et al., Unraveling ancestry, kinship, and violence in a Late Neolithic mass grave. Proc. Natl. Acad. Sci. U. S. A. 116, 10705–10710 (2019).

41. K.-G. Sjögren, et al., Kinship and social organization in Copper Age Europe. A cross-disciplinary analysis of archaeology, DNA, isotopes, and anthropology from two Bell Beaker cemeteries. bioRxiv, 863944 (2019).

42. M. Kircher, S. Sawyer, M. Meyer, Double indexing overcomes inaccuracies in multiplex sequencing on the Illumina platform. Nucleic Acids Res. 40, e3 (2012).

43. M. P. Verdugo, et al., Ancient cattle genomics, origins, and rapid turnover in the Fertile Crescent. Science 365, 173–176 (2019).

44. H. Li, R. Durbin, Fast and accurate short read alignment with Burrows-Wheeler transform. Bioinformatics 25, 1754–1760 (2009).

45. A. Tarasov, A. J. Vilella, E. Cuppen, I. J. Nijman, P. Prins, Sambamba: fast processing of NGS alignment formats. Bioinformatics 31, 2032–2034 (2015).

46. A. McKenna, et al., The Genome Analysis Toolkit: a MapReduce framework for analyzing next-generation DNA sequencing data. Genome Res. 20, 1297–1303 (2010).

47. H. Jónsson, A. Ginolhac, M. Schubert, P. L. F. Johnson, L. Orlando, mapDamage2.0: fast approximate Bayesian estimates of ancient DNA damage parameters. Bioinformatics 29, 1682–1684 (2013).

48. Z. Hofmanová, et al., Early farmers from across Europe directly descended from Neolithic Aegeans. Proc. Natl. Acad. Sci. U. S. A. 113, 6886–6891 (2016).

49. I. Mathieson, et al., Genome-wide patterns of selection in 230 ancient Eurasians. Nature 528, 499–503 (2015).

50. G. D. Poznik, Identifying Y-chromosome haplogroups in arbitrarily large samples of sequenced or genotyped men. bioRxiv (2016) https://doi.org/10.1101/088716.

51. H. Weissensteiner, et al., HaploGrep 2: mitochondrial haplogroup classification in the era of high-throughput sequencing. Nucleic Acids Research 44, W58–W63 (2016).

52. C. Wang, X. Zhan, L. Liang, G. R. Abecasis, X. Lin, Improved Ancestry Estimation for both Genotyping and Sequencing Data using Projection Procrustes Analysis and Genotype Imputation. The American Journal of Human Genetics 96, 926–937 (2015).

53. I. Lazaridis, et al., Genomic insights into the origin of farming in the ancient Near East. Nature 536, 419–424 (2016).

54. N. Patterson, et al., Ancient admixture in human history. Genetics 192, 1065–1093 (2012).

55. M. Lipatov, K. Sanjeev, R. Patro, K. Veeramah, Maximum Likelihood Estimation of Biological Relatedness from Low Coverage Sequencing Data. bioRxiv (2015) https://doi.org/10.1101/023374.

56. 1000 Genomes Project Consortium, et al., An integrated map of genetic variation from 1,092 human genomes. Nature 491, 56–65 (2012).

57. J. M. M. Kuhn, M. Jakobsson, T. Günther, Estimating genetic kin relationships in prehistoric populations. PLOS ONE 13, e0195491 (2018).

58. C. C. Chang, et al., Second-generation PLINK: rising to the challenge of larger and richer datasets. Gigascience 4, 7 (2015).

59. G. Bhatia, N. Patterson, S. Sankararaman, A. L. Price, Estimating and interpreting FST: The impact of rare variants. Genome Research 23, 1514–1521 (2013).

